# Chronic pharmacologic manipulation of dopamine transmission ameliorates metabolic disturbance in syndrome caused by mutated trappc9

**DOI:** 10.1101/2024.02.13.580023

**Authors:** Yan Li, Muhammad Usman, Ellen Sapp, Yuting Ke, Zejian Wang, Adel Boudi, Marian DiFiglia, Xueyi Li

## Abstract

Loss-of-function mutations of the gene encoding the trafficking protein particle complex subunit 9 (trappc9) cause intellectual disability and obesity by unknown mechanisms. Genome-wide analysis links trappc9 to non-alcoholic fatty liver disease (NAFLD). The abrogation of trappc9 in mice has been shown to alter the density of neurons containing dopamine receptor D2 (DRD2) and/or DRD1 in the striatum. Here, we report that trappc9 deficiency in mice resulted in disruption of systemic glucose homeostasis and onset of obesity and NAFLD, which were relieved upon chronic treatment combining DRD2 agonist quinpirole and DRD1 antagonist SCH23390. The homeostasis of systemic glucose in trappc9-deficient mice was restored upon administrating quinpirole alone. Transcriptomic and proteomic analyses revealed signs of impairments in neurotransmitter secretion in trappc9-deficient mice. Brain examinations showed that trappc9-deficient mice synthesized dopamine normally, but their dopamine-secreting neurons had a lower abundance of structures for releasing dopamine in the striatum. Our study suggests that trappc9 loss-of-function causes obesity and NAFLD by constraining dopamine transmission.

## Introduction

Loss-of-function mutations of the gene coding for the trafficking protein particle (TRAPP) complex subunit 9 (trappc9) cause an autosomal recessive syndrome with intellectual disability and malformations of various brain structures; over half of the cases exhibit obesity (*1–11*). Genome-wide analyses detect differential methylation of the trappc9 gene in children with severe obesity as well as in women with a high body-mass index (*12–14*) and link trappc9 to non-alcoholic fatty liver disease (NAFLD) (*15*), the most common liver disease frequently associated with obesity (*16–18*). Consistent with a role for trappc9 in metabolic homeostasis, global metabolic changes are detected in cells from patients with trappc9 mutations (*11*). These human object studies define trappc9 as a risk factor for obesity. Trappc9 is a component specific for the transport protein particle II (TRAPPII) and interacts with trappc10 to form a subcomplex that directs TRAPP towards activating Ypt31/32/Rab11 (*19*), a master player in recycling endocytosed receptors, transporters, and other critical plasma membrane constituents in mammalian cells. Trappc9 is also known as a binding protein for NIK and IKKβ and potentiates the activation of NF-κB (*20*), a signaling pathway that plays a critical role in the pathogenesis of obesity (*21–23*). Exactly how trappc9 variations trigger the onset of obesity and NAFLD is unknown.

We and others recently generated genetically modified mice with loss-of-function mutations in the mouse trappc9 gene; homozygous mutant or knockout (KO) mice of all these 4 mouse lines show deficits in a wide range of behaviors and abnormal changes in various brain structures (*24–27*), replicating what happens in patients. Compared with their wildtype (WT) counterparts, trappc9 KO mice are normal in gaining body weight in early developmental periods but appear overweight shortly after being weaned (*24–27*). Adipose-derived stromal or stem cells (ASCs) play an important role in the development of obesity by remodeling adipose tissues through generating new adipocytes and secreting bioactive molecules to maintain the immunological homeostasis in adipose tissues (*28–31*). ASCs cultured from trappc9 KO mice prefer adipogenic differentiation and contain abnormally large lipid droplets after being induced toward adipogenic differentiation (*32*). Additionally, our trappc9 KO mice at the age period when they start to appear overweight manifest a reduced density of neurons containing dopamine receptor D2 (DRD2) and an increased density of neurons containing DRD1 in the dorsal (caudate/putamen) and ventral (nucleus accumbens, NAc) striatum (*24*). Dopamine transmission plays an important role in body weight control and systemic glucose homeostasis (*33–38*). However, it is unclear whether the incidence of overweight and altered densities of dopamine-receptive neurons in trappc9 KO mice are connected.

In this study, we examined whether trappc9 KO mice were bona fide obese and explored the mechanism likely to be involved. Our studies showed that trappc9 KO mice had global metabolic disturbances and developed NAFLD. Gene expression analysis of ASCs suggested the involvement of dopamine transmission. Chronic manipulation of dopamine transmission counteracted abnormal body weight gain and alleviated lipid accumulation in adipose and liver tissues in trappc9 KO mice. Transcriptomic analysis of DRD2-containing neurons and proteomic analysis of brain synaptosomes pointed to impaired neurotransmitter secretion. Biochemical and histological studies of brain tissues revealed that trappc9 KO mice had a normal capacity in synthesizing dopamine in the brain, but their dopamine-secreting had a reduced abundance of structures for releasing dopamine in the striatum. Our study suggests that trappc9 gene variations trigger the development of obesity and NAFLD by constraining dopamine transmission and provides a potential treatment.

## Results

### Trappc9 deficiency increases body adiposity and disturbs the homeostasis of systemic glucose in mice

Obesity is frequently seen in patients with trappc9 mutations (*11*). Genome-wide analysis detects differential methylation in the trappc9 gene of severely obese children and of women with a high body-mass index (*12–14*). These human studies define trappc9 to be a risk factor for obesity. Consistently, all four trappc9 mutant mouse lines available to date appear overweight postnatally (*24–27*). However, how trappc9 mutations cause obesity is an enigma. Trappc9 KO mice were not different from WT mice in daily intake of water and food (Figure S1), suggesting that other defects are involved. To determine whether trappc9 KO mice were metabolically obese, we first analyzed serum biochemistry profiles. Plasma levels of triglycerides, cholesterol, low-density proteins, high-density proteins, glucose, insulin, prolactin, and leptin were all significantly higher in trappc9 KO mice at age 4 – 5 months than in age-matched WT mice, whereas plasma levels of urea and creatinine were normal (Figure 1A – 1E). Hyperglycemia was detectable in trappc9 KO mice at age 1 month and persisted thereafter (Figure 1F). Glucose and insulin tolerance tests showed that trappc9 KO mice were not different from WT mice in responding to challenges with concentrated glucose and/or insulin (Figure 2G and 2H), suggesting that insulin sensitivity is still retained in C9KO mice at age 4 – 5 months. We then examined the adipose tissues of WT and trappc9 KO mice. The degree of deposition of white fat in the inguinal (iWAT) and gonadal (gWAT) compartments and brown fat (BAT) in the interscapular compartment was increased in trappc9 KO mice relative to age- and sex-matched WT mice (Figure 2A, 2B). Histological examination revealed a decrease in the cell density and an increase in the size (perimeter) of adipocytes in adipose tissues of trappc9 KO mice (Figure 2C – 2E). The adipocytes in brown fat of WT mice were multilocular, whereas those in trappc9 KO mice were predominantly unilocular (Figure 2C). These observations suggest that trappc9 KO mice are bona fide metabolically obese.

**Figure 1.**
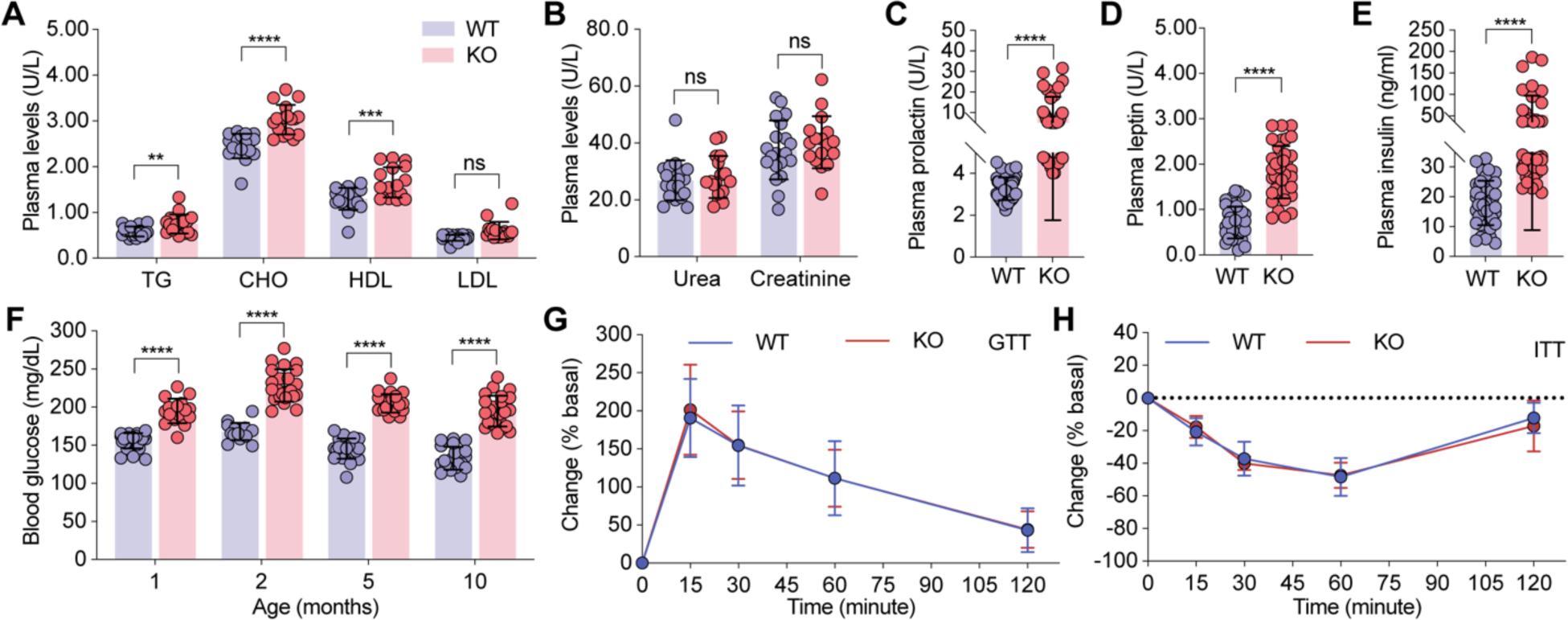
Metabolic disturbance in trappc9-deficient mice. Serum biochemistry analysis shows that lipid profiles **(A**) were elevated whereas levels of urea and creatinine (**B**) were unchanged in trappc9 KO mice relative to age-matched WT mice. Trappc9 KO mice also exhibited hyperprolactinemia (**C**), hyperleptinemia (**D**), hyperinsulinemia (**E**), and hyperglycemia (**F**), which was detectable at age 1 month and persisted thereafter. Glucose (**G**) and insulin (**H**) tolerance tests reveal that trappc9 KO mice responded normally to the challenge with glucose and insulin. Each symbol in bar graphs represents a mouse. N=20 mice (**A**, **B, F**), 30 mice (**C**, **D**, **E**), and 12 mice (**G**, **H**) per genotype. Unless indicated, the age of mice was 5 to 6 months. Data are Mean±SD. Two-tailed Student’s t-test: ** P<0.01; *** P<0.005; **** P<0.001; ns, no significance.

**Figure 2.**
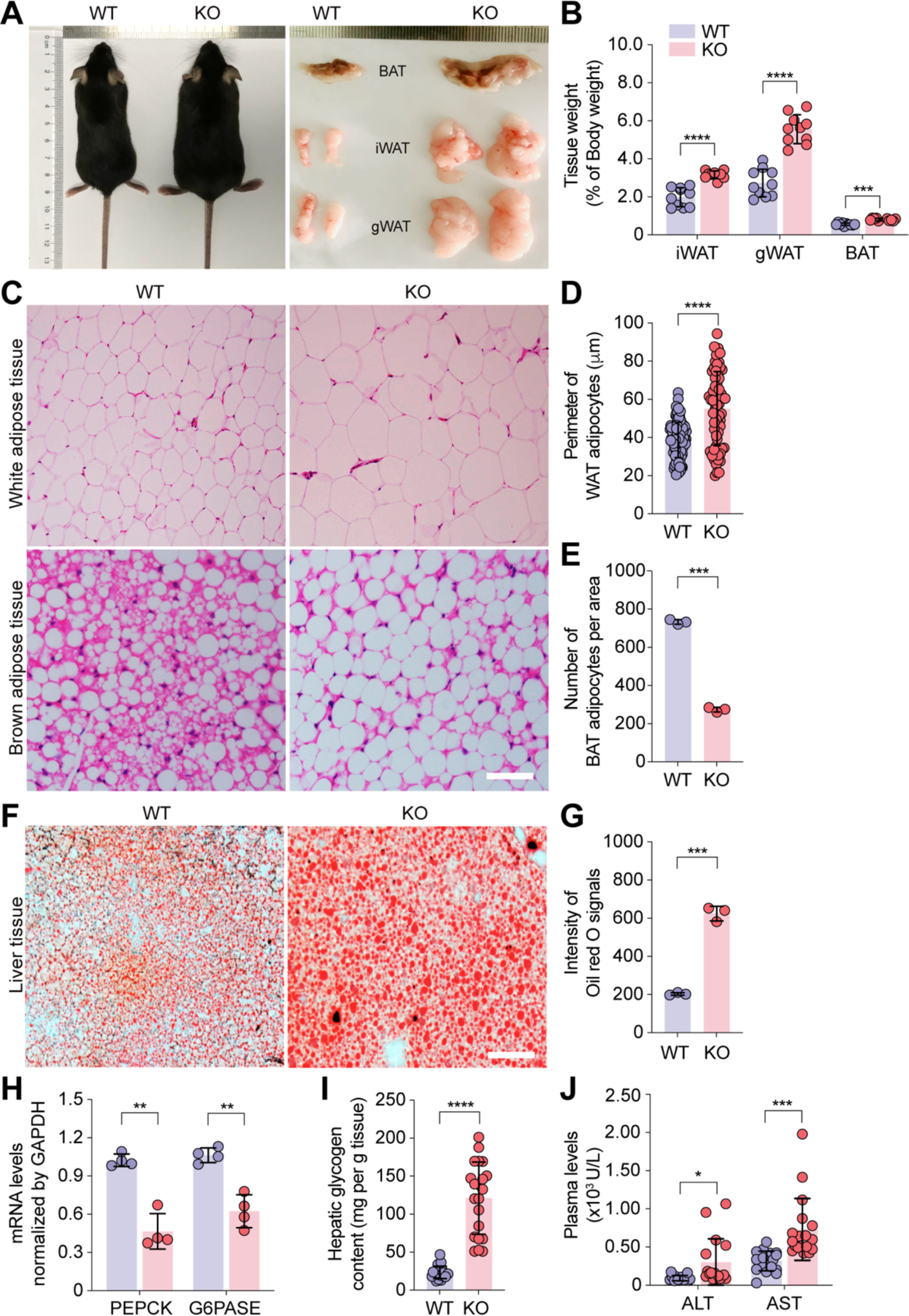
**Trappc9-deficient mice develop obesity and NAFLD. A**) Comparison of trappc9 KO and its littermate, and fat tissues dissected from the inguinal (iWAT), gonadal (gWAT), and interscapular (BAT) compartments. Photographs were taken by Yuting Ke and Muhammad Usman. **B**) Comparison of fat tissue weight in percentage of body weight between trappc9 KO and WT mice. **C**) Hematoxylin and eosin staining of adipose tissues followed by measuring the perimeter (**D**) of adipocytes in white adipose tissues and the number (**E**) of adipocytes in brown adipose tissues. **F**) Oil red O staining of liver tissues followed by densitometric quantification of Oil red O-stained signals (**G**). **H**) Quantitative PCR analysis of gene transcripts for phosphoenolpyruvate carboxykinase (PEPCK) and glucose-6-phosphatase (G6PASE) in the liver, and **I**) measurements of hepatic glycogen contents. **J**) Serum biochemistry analysis of the release of alanine aminotransferase (ALT) and aspartate aminotransferase (AST) into the circulating blood in WT and trappc9 KO mice. Scale bars in (**C**, **F**): 50μm. Each symbol in bar graphs represents one mouse (**B**, **E**, **H**, **I**, **J**) or one cell (**D**). Shown in (**C**, **F**) are representative images from one of 3 mice per genotype. At least 3 sections from different compartments (**C**) or lobes (**F**) were analyzed. The age of mice was 5 to 6 months. Data are Mean±SD. Two-tailed Student’s t-test: * P<0.05; ** P<0.01; *** P<0.005; **** P<0.001.

### Trappc9-deficient mice develop non-alcoholic fatty liver disease

Genome-wide association studies link trappc9 to NAFLD (*15*), a common comorbidity of obesity (*16–18*). Liver enlargement is seen in trappc9 KO mice (*24*). To examine whether there was abnormal accumulation of lipids in the liver, liver tissues from trappc9 KO mice and their WT counterparts were sectioned and subjected to staining with the Oil red O (Figure 2F). Densitometry analysis of Oil red O signals showed that the degree of lipid deposition in hepatocytes of trappc9 KO mice was significantly greater than that in hepatocytes of WT mice (Figure 2G). The liver is the primary organ where excess glucose is stored as glycogen and synthesizes glucose through gluconeogenesis when the level of systemic glucose drops. Quantitative PCR analysis was conducted to examine gene transcripts for glucose-6-phosphatase (G6PASE) and phospho-enolpyruvate carboxykinase (PEPCK), both of which play a critical role in gluconeogenesis, and showed that their transcription was downregulated in the liver of trappc9 KO mice (Figure 2H). On the other hand, the content of hepatic glycogen was elevated in trappc9 KO mice relative to that in WT mice (Figure 2I). These findings imply reprogramming of glucose metabolism in the liver of trappc9 KO mice. The elevated plasma levels of alanine aminotransaminase (ALT) and aspartate aminotransferase (AST) suggest liver function damages in trappc9 KO mice (Figure 2J). Collectively, our results indicate the incidence of NAFLD in trappc9 KO mice.

### Pharmacologic manipulation of dopamine transmission restores the homeostasis of circulating glucose

Having demonstrated that trappc9 KO mice develop obesity and NAFLD, we then looked for the underlying mechanisms. ASCs play an important role in the development of obesity by remodeling adipose tissues through generating new adipocytes and secreting bioactive molecules to maintain the immunological homeostasis in adipose tissues (*28–31*). ASCs cultured from trappc9 KO mice exhibit defects in multiple cellular pathways and are prone to senescence (*32*). To disclose the molecular changes, ASCs were isolated from abdominal fat pads of 1-month old trappc9 KO mice and WT littermates and directly subjected to RNA preparations without in vitro expansion. Bulk RNA-sequencing (RNAseq) analysis identified 15 downregulated and 84 upregulated protein-coding genes in ASCs of trappc9 KO mice (Figure 3A, table S1). Consistent with our previous finding of infertility of male trappc9 KO mice, GO function analysis revealed alterations in reproductive processes and spermatogenesis (Figure 3B, table S2). KEGG pathway analysis suggested disturbances in carbohydrate, lipid, and amino acid metabolism and the involvement of dopaminergic synapse in trappc9 KO mice (Figure 3C, 3D, table S3). Neurotensin receptor (NtsR) 1 and 2 were among the top hits of the identified genes differentially expressed in trappc9 KO ASCs (Figure 3A). STRING functional protein interaction network analysis revealed interplay between the neurotensin and dopamine systems (Figure 3E). The neurotensin system plays a pivotal role in body weight control and regulates dopamine secretion in the brain (*39–42*). These data are consistent with our prior finding that dopamine transmission is altered in trappc9 KO mice (*24*). As the treatment combining DRD1 antagonist SCH23390 and DRD2 agonist quinpirole is effective in ameliorating deficits associated with multiple behaviors in trappc9 KO mice (*24*), we reasoned that this treatment might also be beneficial in controlling obesity and/or NAFLD in trappc9 KO mice.

**Figure 3.**
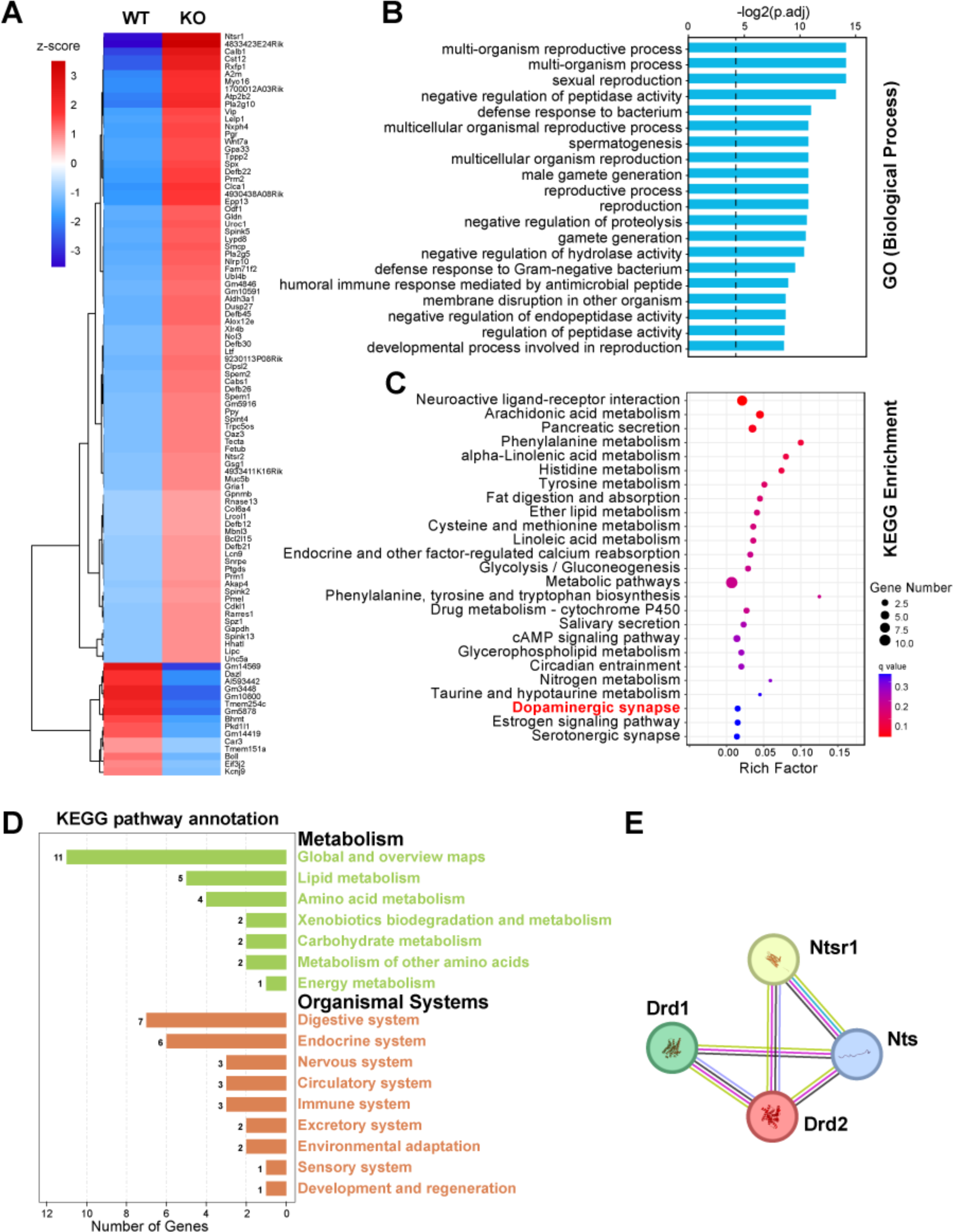
Transcriptomic analysis of adipose-derived stem cells reveals metabolic disturbance and the involvement of dopamine synapse in trappc9-deficient mice. **A**) Heatmap shows differentially expressed genes identified by RNAseq analysis of adipose-derived stem cells in trappc9 KO mice. ASCs were isolated from abdominal fat pads of 1-month old WT and trappc9 KO mice (N=3 to 5 mice per genotype) and directly subjected to RNA preparations without culture in vitro. GO function (**B**) and KEGG pathway (**C**, **D**) analyses of differentially expressed genes in ASCs of trappc9 KO mice. Shown in (**B**, **C**) are top 25 GO biological processes or KEGG pathways. KEGG pathway annotation shows metabolic disturbances in multiple systems (**D**). **E**) The STRING functional protein association network shows interplay between the neurotensin and dopamine systems.

To challenge this idea, we first examined whether the treatment with SCH23390 and quinpirole mitigated hyperglycemia, hyperinsulinemia and hyperprolactinemia in trappc9 KO mice, as dopamine transmission modulates glucose homeostasis and inhibits prolactin and insulin secretion (*43–46*). A cohort of trappc9 KO and WT mice were treated daily by i.p. a dose of SCH23390 and quinpirole. Levels of blood glucose were measured 24 hours after each dose. The treatment had no effect on blood glucose levels in WT mice, but effectively lowered blood glucose of trappc9 KO mice 5 days after the treatment; drug effects lasted for 2 weeks after the treatment was stopped (Figure S2A). The SCH23390 and quinpirole treatment also effectively corrected the hyperinsulinemia and hyperprolactinemia in trappc9 KO mice and resulted in corresponding changes in brain levels of cAMP (Figure S2B – S2D), indicating their actions in the brain. We then assessed whether chronic treatment with both SCH23390 and quinpirole prevented the development of obesity and/or NAFLD in trappc9 KO mice. To this end, three cohorts of trappc9 KO and WT mice, more specifically postnatal week 2, 14 and 46, were designated based on their age when the treatment was started (Figure 4A). The treatment was given daily to mice in the Week-2 cohort for 6 weeks and to mice in both Week-14 and Week- 46 cohorts for 4 weeks (Figure 4A). As expected, the levels of blood glucose in all 3 cohorts of trappc9 KO mice were lowered to normal levels with the treatment (Figure 4B), suggesting that the disturbance of system glucose homeostasis in trappc9 KO mice arises from altered dopamine transmission and is controllable with drugs acting on dopamine receptors.

**Figure 4.**
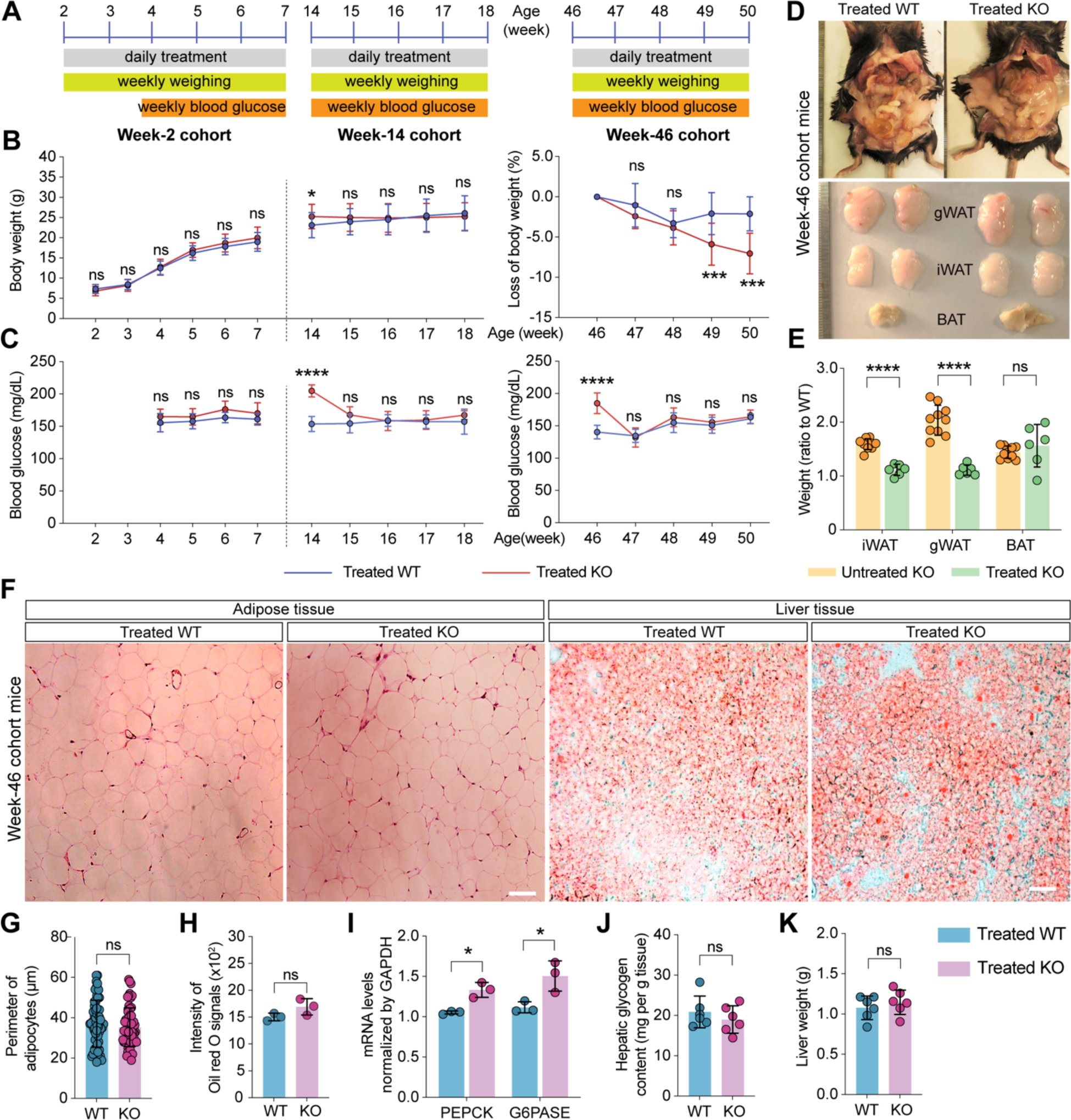
**Chronic pharmacologic manipulation of dopamine transmission alleviates obesity and NAFLD in trappc9-deficient mice. A**) Cohort assignment and experimental schedules. Each cohort contained 16 mice per genotype, 8 per gender. **B**) Body weight (Week-2 and Week-14 cohorts) or body weight loss (Week-46 cohort), and **C**) blood glucose levels in WT and trappc9 KO mice treated daily with SCH23390 and quinpirole for consecutive 6 (Week-2 cohort) or 4 (Week-14 and Week-46 cohorts) weeks. Body weight and blood glucose levels from 16 mice in each cohort at each time point were averaged and graphed. After the drug treatment, mice in the Week-46 cohort were sacrificed for examining body adiposity (**D**, **E**), adipocyte size (**F**, **G**), lipid deposition in liver (**F, H**), PEPCK and G6PASE gene transcripts in liver (**I**), hepatic glycogen contents (**J**), and liver weight (**K**). The adipose tissues were dissected from the corresponding compartments (**D**) and weighed. Photographs in (**D**) were taken by Muhammad Usman and Yan Li. Adipose tissue weight as percentage of body weight was used for calculating the ratio of trappc9 KO adipose tissue weight to WT adipose tissue weight and graphed (**E**). **F**) Images of hematoxylin and eosin-stained adipose tissue sections and Oil red O-stained liver tissue sections. For densitometry, images captured from at least 3 sections from different compartments (adipose tissue) or lobes (liver) for each mouse were analyzed (N=3 mice per genotype). Each symbol in bar graphs represents one mouse (**E**, **H**, **I**, **J**, **K**) or one cell (**G**). Data are Mean±SD. Two-tailed Student’s t-test: * P<0.05; **** P<0.001; ns, no significance.

### The SCH23390 and quinpirole treatment counteracts the abnormal gain of body weight in trappc9-deficient mice

Mice in the Week-2 cohort are in the weaning period. While the body weight of WT mice in the Week-14 cohort is relatively stable, trappc9 KO mice at this age continue to gain body weight (*24*). Hence, the effect of the SCH2339 and quinpirole combined treatment on body weight in trappc9 KO mice in both Week-2 and Week-14 cohorts was determined by directly comparing the body weight of trappc9 KO mice with that of WT mice, whereas body weight loss was used as a measure of drug effects on body weight in the Week-46 cohort. With treatment, trappc9 KO and WT mice in the Week-2 cohort exhibited a similar rate in gaining body weight, and trappc9 KO mice in the Week-14 cohort stopped gaining body weight (Figure 4C). The combined treatment caused trappc9 KO mice in the Week- 46 cohort to lose more weight than their WT counterparts (Figure 4C). These data suggest that pharmacologic manipulation of dopamine signaling counteracts the abnormal gain of body weight in trappc9-deficient mice.

### The SCH23390 and quinpirole combined treatment alleviates lipid accumulation in adipose and liver tissues in trappc9-deficient mice

To examine the effect of the combined treatment on body adiposity, mice from the Week- 46 cohort were sacrificed after treatment. The adipose tissues from abdominal, gonadal, and inguinal compartments as well as the liver of each sacrificed mouse were harvested for examination. The abdominal fat deposition in trappc9 KO mice was not overtly different from that in WT mice after the combined treatment for 4 weeks (Figure 4D). The inguinal (iWAT) and gonadal (gWAT) as well as interscapular (BAT) fat depots from drug-treated trappc9 KO mice appeared similar in size (Figure 4D). A comparison of the ratio of trappc9 KO adipose tissue weight (as percentage of body weight) to WT adipose tissue weight (as percentage of body weight) between drug-treated trappc9 KO mice and untreated trappc9 KO mice showed that the combined treatment resulted in a significant reduction of white adipose tissues (Figure 4E). Histological examination of white adipose tissues revealed that the size (perimeter) of adipocytes in drug-treated trappc9 KO mice was no longer different from that in drug-treated WT mice (Figure 4F, 4G), suggesting that lipid accumulation in adipose tissues was depressed with the combined treatment. In summary, the SCH23390 and quinpirole combined treatment effectively relieves body adiposity in trappc9 KO mice.

Upon the combined treatment, Oil red O signal in liver sections of trappc9 KO mice was not different from that in liver sections of WT mice receiving the same treatment (Figure 4F, 4H), indicating that the treatment attenuates lipid accumulation in the liver of trappc9 KO mice. Quantitative PCR analysis found that the levels of transcripts for G6PASE and PEPCK in the liver of trappc9 KO mice were reversed (Figure 4I). Concomitantly, the content of hepatic glycogen in drug-treated trappc9 KO mice was lowered to normal levels (Figure 4J). These results suggest that the combined treatment restores glucose metabolism in the liver of trappc9 KO mice. The weight of the liver of trappc9 KO mice receded to the level of WT mice (Figure 4K). Altogether, our data suggest that chronic treatment with drugs acting on dopamine receptors blunts the development of obesity and NAFLD in trappc9-deficient mice.

### Deficient stimulation of DRD2-containing neurons underlies the homeostatic disturbance systemic glucose in trappc9-deficient mice

Our above studies showed that the treatment combining SCH23390 and quinpirole restored the homeostasis of systemic glucose and counteracted obesity and NAFLD. To determine whether the disruption of systemic glucose homeostasis was primarily driven by altered dopamine transmission through DRD1 or DRD2 or both, a cohort of mice was treated with either quinpirole or SCH23390. Our results showed that quinpirole effectively reduced blood glucose of trappc9 KO mice (Figure 5A), yet SCH23390 had a trivial effect on blood glucose levels in trappc9 KO mice (Figure 5B), suggesting that deficient stimulation of DRD2 is the key contributor. As DRD2 is also expressed in peripheral tissues, e.g., the pancreas and adipose tissues (*47–49*), we determined whether the blood glucose-lowering effect of quinpirole in trappc9 KO mice was achieved through its central or peripheral actions. To this end, we gave mice daily by i.p. a dose of dopamine, which is unable to pass through the blood-brain-barrier and thus can act only in the peripheral tissues. The idea was that if quinpirole lowered blood glucose in trappc9 KO mice through its peripheral actions, dopamine should also be effective. However, the level of blood glucose in trappc9 KO mice did not decline upon daily treatment with dopamine over 8 days (Figure 5C), indicating that the disturbance of systemic glucose homeostasis most likely stems from deficient stimulation of DRD2 in the brain. To examine whether the suppression of DRD1 was necessary for dopamine to lower blood glucose in trappc9 KO mice, mice were treated daily with SCH23390 combined with dopamine. Our data showed that SCH23390 combined with dopamine did not change blood glucose levels in trappc9 KO mice as well (Figure 5D). Collectively, these data suggest that the disruption of systemic glucose homeostasis in trappc9 KO is a consequence of deficient stimulation of DRD2 in the brain.

**Figure 5.**
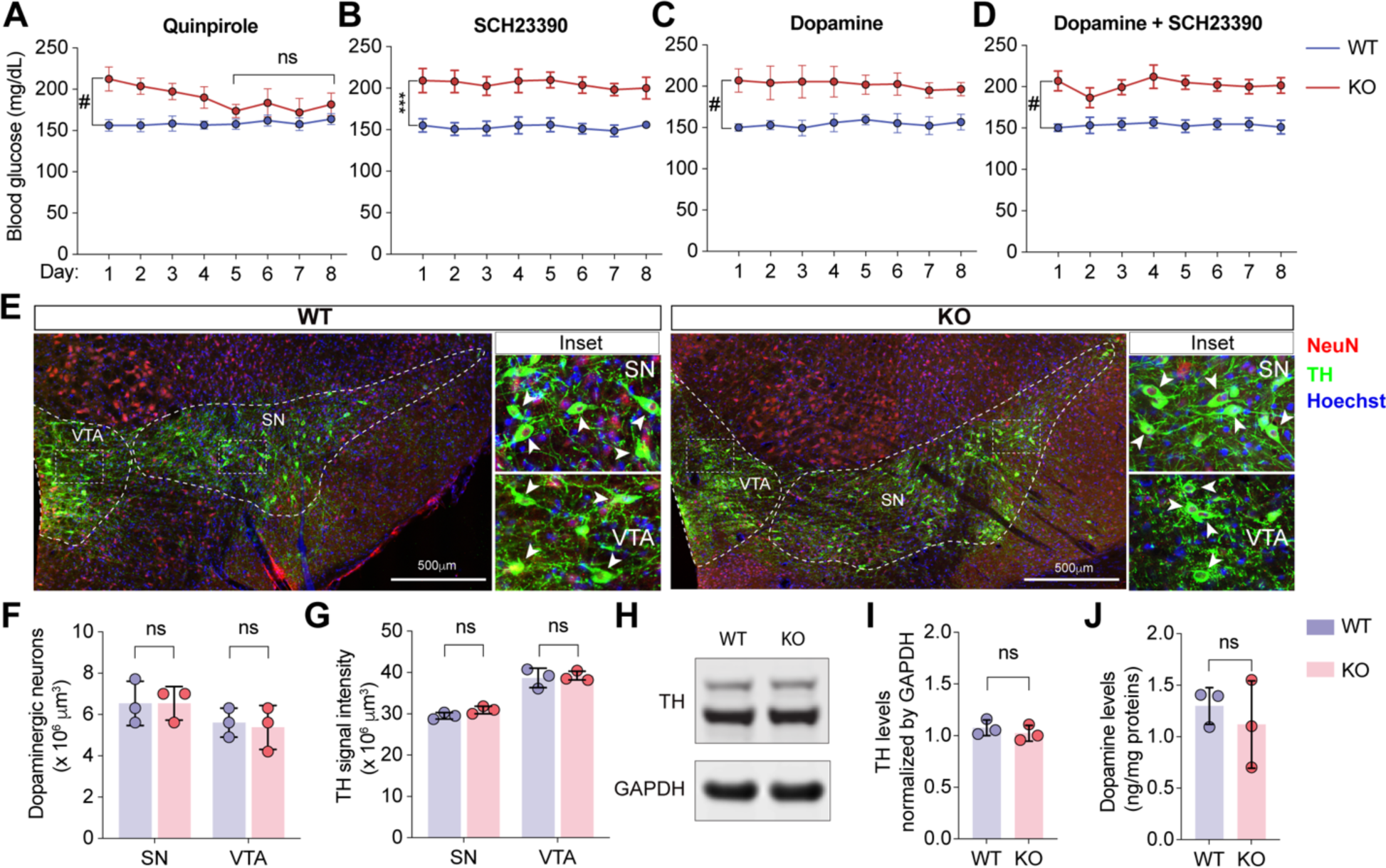
Trappc9 deficiency disrupts systemic glucose homeostasis by abating D2 receptor stimulation without affecting dopamine synthesis in the brain. Comparisons of blood glucose levels in WT and trappc9 KO mice receiving daily treatment, by i.p., with quinpirole alone for 5 days (**A**) or with SCH23390 alone (**B**) or with brain-impermeable dopamine (**C**) or with dopamine combined with SCH23390 (**D**) for consecutive 8 days. **E**) Confocal images taken from the midbrain area of WT and trappc9 KO mouse brain sections stained with antibodies for neuronal marker NeuN (red), dopaminergic neuronal marker TH (green), and DNA dyes (blue). Boxed regions within the VTA and SN, respectively, were enlarged as Inset. VTA, ventral tegmental area; SN, substantia nigra. Stereology counting of somata containing both NeuN and TH (**F**) and densitometric quantification of TH signals (**G**) within the indicated area closed with a contour in confocal images (**E**). Confocal images used for quantitative analyses were obtained from 3 consecutive brain sections from each brain (N=3 mice per genotype). **H**) Western blot analysis of whole brain lysates followed by densitometry of TH immunoreactive signals (**I**). **J**) Comparison of dopamine levels in whole brain lysates of WT and trappc9 KO mice. Pharmacological studies in (**A**, **B**, **C**, **D**) were performed with 7 mice per genotype. Each symbol in bar graphs (**F**, **G**, **I**, **J**) represents one animal. Data are Mean±SD. Two-tailed Student’s t-test: *** P<0.005; # P<0.0001; ns, no significance.

### Trappc9-deficient mice are normal in synthesizing dopamine but exhibit signs of impairments in dopamine secretion

The effectiveness of SCH23390 and quinpirole in relieving obesity and NAFLD in trappc9 KO mice indicates that when ligands are sufficiently available, DRD1- and/or DRD2- containing neurons as well as their effector cells in trappc9 KO mice function normally. Hence, we examined whether there was a loss of capacity of synthesizing dopamine in trappc9 KO mice. The dopaminergic neurons in the midbrain (the substantia nigra pars compacta and the ventral tegmental area) are the major sources of dopamine in the brain (*49, 50*). Histological and Western blot analyses showed that compared with WT mice, trappc9 KO contained a normal cell density of dopaminergic neurons in the midbrain and expressed comparable levels of tyrosine hydroxylase (TH), the rate-limiting enzyme in dopamine synthesis (Figure 5E – 5I). Consistently, similar levels of dopamine were detected in brain lysates of trappc9 KO and WT mice (Figure 5J). These results suggest that trappc9 KO mice synthesize dopamine normally in the brain.

The above studies suggest that deficient stimulation of DRD2 in the brain was a key contributor to the disruption of systemic glucose homeostasis in trappc9 KO mice. To disclose potential molecular changes, DRD2-containing neurons were immunoisolated from trappc9 KO mice and WT littermates and processed for RNAseq analysis (Figure 6A). DRD2 transcript was detected but not altered in trappc9 KO mice (table S4). However, our analysis identified 6 downregulated and 37 upregulated protein-coding genes in DRD2-containing cells of trappc9 KO mice (Figure 6B, table S4). GO and KEGG analyses implied alterations in neurotransmitter secretion and synapse formation in trappc9 KO mice (Figure 6C, table S5 and S6). Among the identified genes, Unc13c (Munc13-3) and Rims1 (RIM) are known to manage dopamine secretion, whereas Gpm6a, Synpo, Insyn1, and Dlg2 (Psd93) are involved in forming and/or stabilizing synapses. NtsR2 was also the top hit of the upregulated genes in DRD2-containing neurons of trappc9 KO mice. Given their functions in synaptic transmission, their protein levels were examined by Western blot analysis. The results showed that the levels of Unc13c, Synpo1, Dlg2, Gpm6a, and Insyn1 were reduced in brain lysates of female trappc9 KO mice relative to their matched WT counterparts (Figure 6D, 6E). While Dlg2 and Gpm6a were also decreased in male trappc9 KO mice, the levels of Unc13c, Synpo1, and Insyn1 were comparable relative to their levels in male WT mice (Figure 6D, 6E). It is not clear whether this divergence is related to the sex difference in time of onset of obesity and severity of phenotypes between female and male trappc9 KO mice (*24, 27*). Notably, the trend of change in protein levels was opposite to that in gene transcripts, likely because compensatory effects have been developed in trappc9 KO mice.

**Figure 6.**
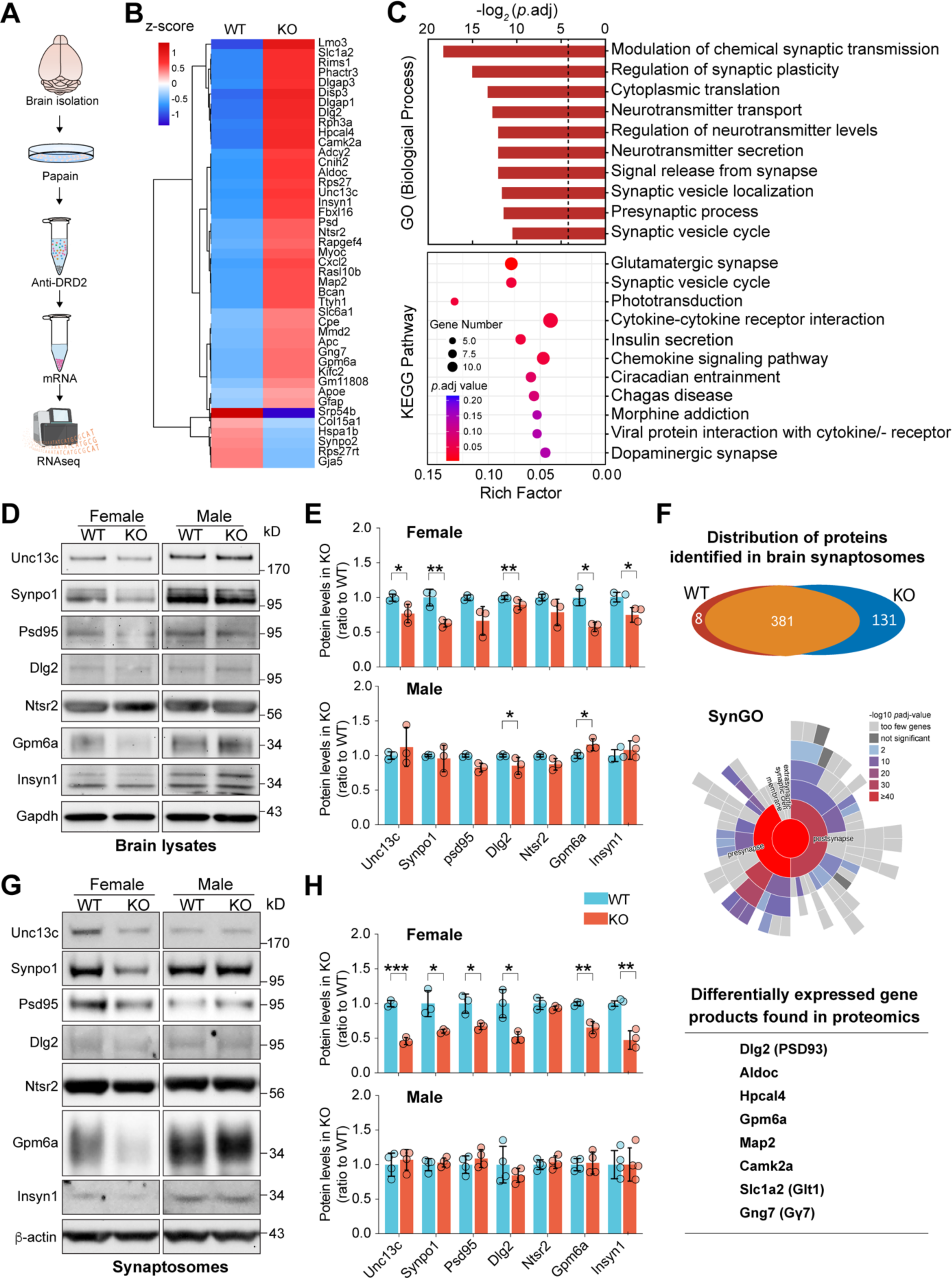
Transcriptomic and proteomic analyses reveal signs of impairments in neurotransmitter secretion in trappc9-deficient mice. A) Experimental schedule for RNAseq analysis of DRD2-containing cells from WT and trappc9 KO mice (N=3 mice per genotype). **B)** Heatmap representation of the differentially expressed genes identified by RNAseq analysis of DRD2-containing cells. **C**) GO function and KEGG pathway analyses of differentially expressed genes. Shown are top 10 GO biological processes and KEGG pathways, respectively. **D**) Western blot analysis of gene products of some differentially expressed genes in brain lysates followed by densitometric quantification of the protein levels (**E**). **F**) Overall distribution of proteins identified by proteomic analysis of brain synaptosomes (Upper panel), Synaptic Gene Ontologies (SynGO) analysis of differentially expressed proteins with a change by at least twofold (Middle panel), and summary of differentially expressed genes identified in both RNAseq and proteomic analyses (Lower panel). **G**) Western blot analysis of the levels of the chosen proteins as in (**D**) in brain synaptosomes of WT and trappc9 KO mice followed by (**H**) densitometric quantification of the protein levels. Each symbol in bar graphs represents one mouse. Data are Mean±SD. Two-tailed Student’s t-test: * P<0.05; ** P<0.01; *** P<0.005.

To further demonstrate synaptic perturbation in trappc9 KO mice, synaptosomes were prepared from fresh brain tissues of 3 – 4 months old WT and trappc9 KO mice. Quantitative proteomics analysis identified 8 proteins only present in WT mouse synaptosomes and 131 were only present in synaptosomes of trappc9 KO mice (Figure 6F upper panel, table S7). Synaptic Gene Ontologies (SynGO) analysis of the differentially expressed proteins with a change by at least twofold showed that both presynaptic and postsynaptic proteins were altered in trappc9 KO mice (Figure 6F middle panel, table S8). It is worth noting that 8 genes/proteins were identified in both RNAseq and proteomic analyses (Figure 6F lower panel). Western blot analysis of the above set of proteins revealed that their levels were diminished in brain synaptosomes from female trappc9 KO mice and unchanged in synaptosomes from male trappc9 KO mice (Figure 6G, 6H). Collectively, these data suggest that the deficiency of trappc9 indeed constrains synapse formation.

### Trappc9 deficiency reduces the density of dopamine axonal varicosities

Having found clues of impairments in neurotransmitter secretion and synapse formation, we examined whether dopamine synapses were altered in trappc9 KO mice. In projection areas, e.g., the striatum, dopaminergic neurons release dopamine at their axonal varicosities, which synapse onto dendrites and/or dendritic spines of dopamine-receptive neurons (*51, 52*). In this regard, dopamine synapse alterations can be reflected from a change in dopamine-secreting axonal varicosities. The abrogation of trappc9 in mice diminishes dendritic spines (*26*) and alters the density of dopamine-receptive neurons (*24*). As dopamine neurotransmission in the NAc is involved in body weight control and regulates systemic glucose homeostasis (*34, 45, 46, 53–55*), we focused our analysis on dopamine axonal varicosities in the NAc. To this end, brain sections containing the striatum were co-labeled with antibodies for TH, active zone scaffold protein Bassoon, and synaptic vesicle-associated protein Vamp2 to identify dopamine-secreting axonal varicosities (Figure 7A). While there was no difference in the frequency of structures positive for Bassoon or for Vamp2 between WT and trappc9 KO mice (Figure 7B, 7C), the structures containing both Bassoon and Vamp2 (total neurotransmitter release sites) were less abundant in the striatum of trappc9 KO than in the striatum of WT mice (Figure 7D), suggesting that trappc9 deficiency reduces the abundance of synapses. Not surprisingly, the frequency of structures positive for both Bassoon and Vamp2 in TH- positive processes was reduced in trappc9 KO mice (Figure 7E). Further analysis revealed that the magnitude of decrease of dopamine-secreting varicosities was significantly greater than that of total neurotransmitter-releasing sites (Figure 7F), suggestive of differential vulnerability of dopamine synapse to the loss of trappc9.

**Figure 7.**
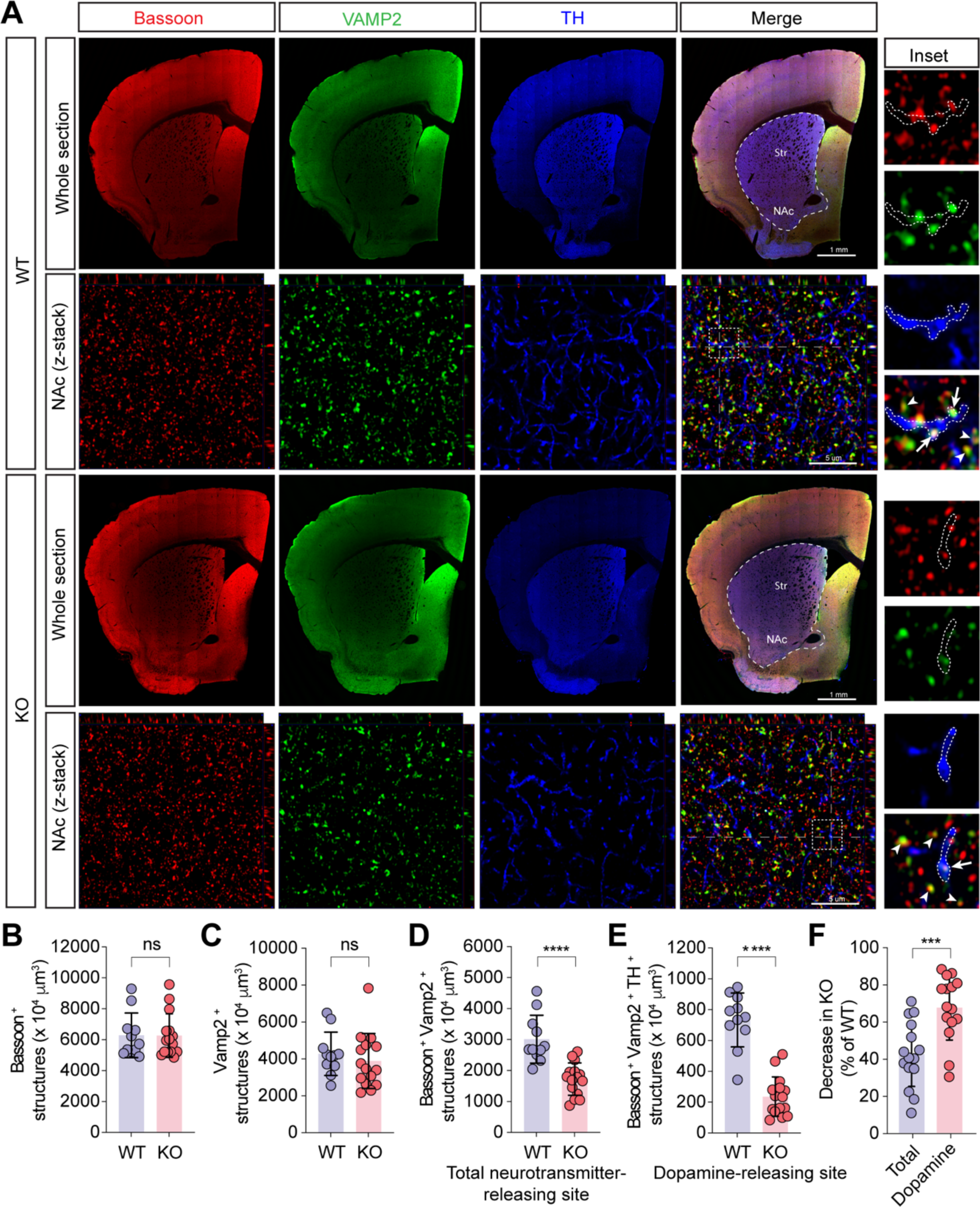
The abundance of dopamine release sites is diminished in the brain of trappc9-deficient mice. A series of 3 consecutive sections per brain (N=3 mice/genotype) were labeled with antibodies for Bassoon, Vamp2, and TH to detect dopamine release sites. **A)** Shown were low-magnification images of a brain section and high-magnification z-stack images taken within the NAc. Boxed regions in the high-magnification merged z-stack images were enlarged as Inset. Z-stack images were used for counting the frequency of structures positive for Bassoon (**B**), or Vamp2 (**C**), or for both Bassoon and Vamp2 (**D**), and the frequency of those Bassoon+ and Vamp2+ structures in TH+ processes (**E**) with stereology methods. The percentage of reduction of structures positive for both Bassoon and Vamp2 (total release sites) and the percentage of reduction of Bassoon+ and Vamp2+ structures within TH+ processes (dopamine release sites) were calculated and graphed (**F**). In enlarged merged images (Inset), arrows point to dopamine release sites, whereas arrowheads indicate the release sites of other neurotransmitters. Contours indicate TH+ axonal segments. Each symbol in bar graphs represents one mouse. Data are Mean±SD. Two-tailed Student’s t-test: *** P<0.005; **** P<0.0001; ns, no significance.

## Discussion

Human object studies have suggested trappc9 to be a risk factor for obesity and NAFLD, but the relevant mechanism is not clear. In this study, we found that the ablation of trappc9 resulted in disruption of systemic glucose homeostasis, which was associated with hyperinsulinemia, hyperleptinemia, hyperprolactinemia, dyslipidemia, adipocyte hypertrophy, and glucose metabolism reprogramming and lipid accumulation in the liver. Transcriptomic analysis of adipose-derived stem cells implied global metabolic changes and the involvement of dopamine transmission. Chronic pharmacologic manipulation of dopamine transmission restored systemic glucose homeostasis and relieved obesity and NAFLD in trappc9 KO mice. We further showed that the disruption of systemic glucose homeostasis in trappc9 KO mice originated from deficient stimulation of DRD2. Transcriptomic and proteomic analyses pointed to impairments in neurotransmitter secretion in trappc9 KO mice. Consistently, trappc9 KO mice synthesized dopamine normally in the brain, but their dopamine-secreting neurons had a reduced abundance of structures for releasing dopamine in the projection areas, e.g., the nucleus accumbens.

Our study suggests that trappc9 loss-of-function causes obesity and NAFLD by perturbing dopaminergic synapse formation or stability.

Along with prior findings that trappc9 ablation compromises rab11 activation and reduces dendritic spines (*19, 24, 26*), our studies support a model for the onset of obesity triggered by the deficiency of trappc9. In this model (Figure 8), impaired activation of rab11 is the driving force. 1) A loss of trappc9 abolishes the formation of fully functional transport protein particle II for activating rab11 and results in rab11 functional decline; 2) a reduction in rab11 function impedes the recycling of internalized plasma membrane lipids and proteins from recycling endosomes (RE) to cell surfaces, thereby disturbing cell functions; 3) chronic impairment of rab11-regulated trafficking diminishes the abundance of dopaminergic synapses; 4) altered dopamine transmission disrupts systemic glucose homeostasis, thereby triggering a cascade of metabolic changes, including excessive secretion of insulin, prolactin and leptin, and lipid deposition in adipose and liver tissues, eventually leading to the onset of obesity and NAFLD.

**Figure 8.**
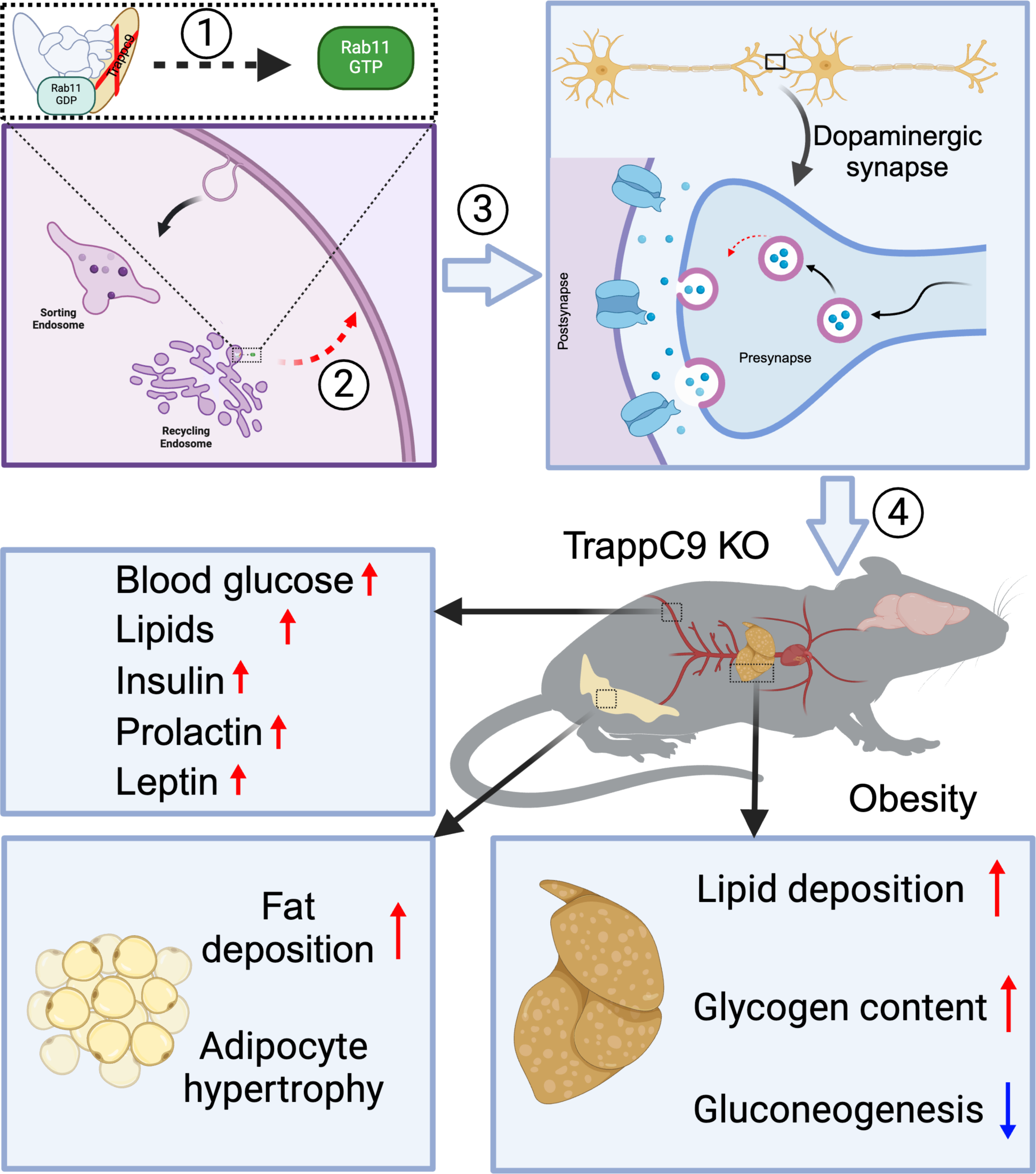
**Schematic representation of obesity onset induced by the deficiency of trappc9**. In this mode, the impaired activation of rab11 is the driving force. 1) The loss of trappc9 abolishes the formation of fully functional transport protein particle II for activating rab11 and results in rab11 functional decline; 2) a reduction in rab11 function impedes endocytic recycling of internalized plasma membrane lipids and proteins from the recycling endosome (RE) to the cell surface, thereby disturbing cell functions; 3) chronic impairment of rab11-based trafficking leads to a decline in the formation of dopaminergic synapses, thereby restraining dopamine transmission; 4) dopamine neurotransmission alteration disrupts systemic glucose homeostasis and triggers a cascade of metabolic changes, including excessive secretion of insulin, prolactin and leptin, and lipid deposition in adipose and liver tissues, and eventually leads to the onset of obesity and NAFLD. The scheme was drawn with BioRender (www.biorender.com).

While the trappc9-caused syndrome is extremely rare, trappc9 variations may contribute significantly to the prevalence of obesity based on the following features of trappc9 pathogenic mutations. Firstly, they are highly diverse, which range from a single nucleotide exchange to indels. Secondly, they are not clustered within certain “hot spots” but scattered throughout the gene and occur in exons, introns, and/or at splicing sites, suggesting that they are not restricted to persons of certain specific geographical regions or races. Thirdly, trappc9 pathogenic mutations can be inherited from parents with no family history of consanguineous marriage (*9, 56*) and even de novo (*57, 58*). Moreover, abnormal epigenetic modification of the trappc9 gene is associated with obesity in humans (*12–14*). Although it is not clear how common trappc9 mutations are in the general population, more than 4, 000 trappc9 variants are included in the gnomAD V4.0 dataset (https://gnomad.broadinstitute.org/gene/ENSG00000167632?dataset=gnomad_r4), in which several known pathogenic mutations are absent.

In mice, trappc9 is imprinted in the brain with a maternal allele-biased expression (∼70%) and shows an equal biallelic expression in peripheral tissues (*25, 59*). Trappc9 mice with only the maternal allele mutated appear overweight, whereas mice with the paternal allele mutated are normal (*25*), indicating that obesity onset in trappc9 KO mice involves a loss of function mainly in the brain. Prior studies reveal that the abrogation of trappc9 diminishes DRD2-containing neurons and increases DRD1-containing neurons in the striatum; acute treatment combining quinpirole and SCH23390 to stimulate DRD2 and repress DRD1 effectively attenuates defects associated with multiple behaviors (*24*). In this study, we found that chronic treatment combining quinpirole and SCH2330 counteracted abnormal body weight gain and alleviated lipid accumulation in adipose and liver tissues in trappc9 KO mice. Quinpirole alone was sufficient to restore blood glucose homeostasis in trappc9 KO mice, but peripheral administration of brain-impermeable dopamine or its combination with SCH23390 had a minor effect. These findings suggest that the development of obesity in trappc9 KO mice is related to an improper function of the dopamine system in the brain.

Dopamine transmission depends on the adequacy of dopamine in extracellular spaces and dopamine receptors on the cell surface. On the cell surface of striatal neurons, levels of DRD1 are increased whereas levels of DRD2 are decreased in trappc9 KO mice (*24*). If dopamine was sufficiently accessible, such changes in DRD1/2 levels on neuronal surfaces should render trappc9 KO mice hyperactive based on the classical view of motor control (*60, 61*). On the contrary, they are hypoactive (*24*). The hypoactivity of trappc9 KO mice is unlikely to stem from insufficient excitatory inputs because animal immobility appears in trappc9 KO mice as quickly as in WT mice after administrating quinpirole or SCH23390 (*24*). As dopamine can bind to its receptors only when it is released into extracellular spaces, altered dopamine transmission in trappc9 KO mice most likely results from defective secretion of dopamine. In projection areas, e.g., the striatum, dopaminergic neurons secret dopamine at axonal varicosities, which synapse onto dendrites and/or dendritic spines of their innervated neurons (*51, 52*). Our studies showed that dopaminergic neurons in trappc9 KO mice had a lower abundance of dopamine release sites on their projection axons in the NAc. This could be related to the decreased expression of proteins involved in neurotransmitter secretion. Alternatively, the decrease of dopamine-releasing sites is a compensatory response to changes in dendritic spines, which are reduced in trappc9 KO mice (*26*). Consistently, we found that proteins related to spine formation, e.g., Gpm6a and Synpo1, were diminished in trappc9 KO mice.

Dopamine impacts systemic glucose homeostasis and body weight through both central and peripheral actions. Centrally, dopamine regulates food intake, thermogenesis, and hormone secretion in the pituitary gland (*37*), whereas peripherally, it modulates the secretin of insulin and adipokines as well as insulin-stimulated uptake of glucose (*44, 62–64*). In this regard, the effect of SCH23390 and quinpirole on glucose metabolism and obesity onset in trappc9 KO mice might be achieved through actions in peripheral tissues. If so, peripheral administration of dopamine combined with SCH23390 should mimic the combination of SCH23390 and quinpirole, because peripherally administrated dopamine is unable to cross the blood-brain barrier and can act only in peripheral tissues. However, neither dopamine nor SCH23390 nor their combination was able to lower blood glucose levels in trappc9 KO mice. On the other hand, quinpirole alone effectively downgraded blood glucose in trappc9 KO mice. Hence, deficient stimulation of DRD2 in the brain is a key contributor.

ASCs are key regulators of adipose tissue remodeling by giving rise to adipocytes and maintaining immunological homeostasis in adipose tissues (*29, 31*). ASC dysfunction and obesity are intertwined to magnify the development of obesity and comorbidities (*31*). Trappc9 KO ASCs prefer adipogenic differentiation and contain abnormally large lipid droplets when being induced to differentiate (*32*). Consistent with these findings, RNAseq analysis of ASCs detected changes in genes related to the metabolism of carbohydrates, lipids, and amino acids in trappc9 KO mice. Hence, ASC dysfunction in trappc9 KO mice may play a causative role in the development of obesity. DRD1 and DRD2 transcripts were detected in ASCs, but their levels were not altered in trappc9 KO ASCs. It is not clear whether and how significantly dopamine signaling is altered in trappc9 KO ASCs.

In addition to trappc9, several other components of TRAPPII are also linked to human diseases, which include trappc2, trappc2L, trappc4, trapp6A/B, and trappc10. In TRAPPII, trappc9 and trappc10 form a subcomplex (*19*); their deficiency mutually depletes each other in mice (*24, 65*). Like trappc9 KO mice, trappc10-deficient mice develop obesity postnatally (*65*). However, patients bearing mutations of trappc2, trappc2L, trappc4, trappc6A/B or trappc10 do not develop obesity as patients with trappc9 mutations. In this regard, obesity onset caused by trappc9 mutations is likely to involve a function independent of TRAPPII. Trappc9 has been shown to potentiate the activation of NF-κB (*20*). Yet, no sign of defective activation of NF-κB was found in trappc9 KO mice (*24, 26*). As deficient NF-κB activation in fibroblasts of patients with trappc9 mutations is seen upon stimulation with tumor necrosis factor alpha (*3*), NF-κB activation may be altered in trappc9 KO mice under induction conditions. Based on these observations, trappc9 deficiency should hamper the onset of obesity, as blunting NF-κB activation abates obesity development (*22, 23*). On the contrary, trappc9 deficiency renders obesity in both humans and mice. Alternatively, NF-κB activation might be enhanced in trappc9 KO mice under stimulation conditions. If so, NF-κB activation in the trappc9 KO setting should occur through another mechanism. DRD2-mediated signaling through β-arrestin 2, Akt, and protein phosphatase 2A represses NF-κB activation (*66*). Future studies are needed to challenge this idea.

## Conclusions

Collectively, this study shows that trappc9 loss-of-function disrupts systemic glucose homeostasis and leads to the onset of obesity and fatty liver disease by reducing dopaminergic synapses, thus establishing the trappc9 mutant mouse line as a tool for studying obesity and non-alcoholic fatty liver disease. Our study provides a potential treatment for obesity and non-alcoholic fatty liver disease.

## Materials and Methods

### Animals

Animals were housed in male and female groups and maintained in a pathogen-free environment under regular 12-hour light/dark cycle with food and water ad libitum. All experimental procedures were carried out following the NIH guide for the Care and Use of Laboratory Animals and approved by the Institutional Animal Care and Use Committee of the Shanghai Jiao Tong University (# A2017029) and/or the Massachusetts General Hospital (#2004N000248). All mice were genotyped by PCR as described (*24*).

### Serum biochemistry profile

Blood samples were collected from WT and trappc9 KO mice and centrifugated at 4°C 3, 000rpm for 12 minutes. The resulting plasmas were subjected to measuring levels of triglyceride, cholesterol, high-density lipoprotein, low-density lipoprotein, urea, creatinine aspartate aminotransferase, and alanine transaminase (Servicebio company, Shanghai).

### Glucose and Insulin Tolerance Tests

Levels of blood glucose were measured with a glucometer (Accu-Check Active, Roche, Germany) following the supplier’s procedures. The first drop of blood was discarded, and the second drop was placed on a test strip, which was then inserted into the glucometer. For the glucose tolerance test (GTT), mice were fasted overnight (14 hours) with access only to drinking water throughout the test. The body weight and fasting blood glucose levels of each mouse were measured and recorded before starting the GTT. Each mouse was given by intraperitoneal injection 2g glucose per kg body weight, which was prepared with 20% glucose (Sigma-Aldrich). Levels of blood glucose were measured at 15, 30, 60, and 120 minutes after concentrated glucose was injected. Blood loss was minimized by slightly pressing and cleaning the incision area with 75% alcohol swabs after each measurement. At the end of the GTT, each mouse was provided with abundant food and water and observed for 1-2 days for any odd behavior. For the insulin tolerance test (ITT), mice were fasted for 6 hours between 7:00 AM and 1:00 PM as above. After fasting, the body weight and levels of blood glucose of each mouse were measured. Each mouse was the given by intraperitoneal injection 0.75IU insulin per kg body weight. Levels of blood glucose were measured at 15, 30, 60, and 120 minutes after insulin was given. After the ITT was completed, mice were transferred to clean cages with free access to food and water and monitored for 2-4 days for abnormal behaviors.

### Enzyme-linked Immunosorbent Assay

Plasma levels of leptin, prolactin, and insulin were measured by ELISA (enzyme-linked immunosorbent assay) with reagents obtained from Ray biotech, CusaBio, and Crystal Chem, respectively, according to the reagent supplier’s instructions. The optical density at 450nm (OD450) of each sample was measured in a multifunctional microplate reader (Biotech, Synergy LX). The absorbance for each set of standards was measured. The standard curve was drawn using the curve expert and GraphPad software, and the best fit curve was drawn through these points.

### Pharmacological treatment

To assess the effect of the combined treatment with SCH23390 and quinpirole on blood glucose, a cohort of WT and trappc9 KO mice at age 3 – 4 months were given daily by intraperitoneal injection (i.p.) a dose of both SCH23390 at 0.1 mg/kg (Tocris, Cat. No. 0925) and quinpirole at 1 mg/kg (Tocris, Cat. No. 1061). Levels of blood glucose were measured 24 hours after each dose. Another cohort of mice at age 3 to 4 months were used for determining whether the blood glucose lowering effect originated from SCH23390, or quinpirole, or both and for determining dopamine at 0.1 mg/kg (dopamine hydrochloride, Sigma-Aldrich, Cat. No. EY1491) in lowering blood glucose levels. For evaluating the effect of the chronic treatment combining SCH23390 (0.1 mg/kg) and quinpirole (1 mg/kg) on obesity and NAFLD, three cohorts of WT and trappc9 KO mice were devised based on their postnatal age when the treatment was started: 2 weeks, 14 weeks, and 46 weeks. The drugs were administered by i.p. every day. Body weight and blood glucose levels were measured weekly.

### Histological examination of adipose tissues

Abdominal white and inter-scapular brown fat tissues were surgically removed from each deeply anesthetized mouse, transferred to a glass vial containing 4% paraformaldehyde and fixed overnight at 4 °C. Fixed adipose tissues were washed several times in PBS, embedded in paraffin and cut into 5 μm sections, which were collected onto glass slides. The sections were then stained in the hematoxylin solution for 5 minutes. After washes in distilled water, the sections on slides were soaked in the eosin reagent for 1-2 minutes and then rinsed in distilled water. The sections were dehydrated sequentially in 75%, 80%, 90%, 95%, and 100% alcohol, 2 minutes in each, and then cleansed twice in xylene, each for 2 minutes. After being dried, the sections were coverslipped with mounting media and examined under an Olympus BX53 microscope. Digital images were taken from at least three randomly chosen visual fields for each section.

### Oil Red O staining of liver tissues

Fresh liver tissues were surgically removed from deeply anesthetized mice, washed multiple times with phosphate buffer saline (PBS), and embedded in OCT media. A series of 15 μm sections were cut on a cryostat (Leica). The sections were directly mounted onto glass slides and stained with 0.5% of Oil red O solution for 10 minutes. The sections were then rinsed with running distilled water for 15 minutes, dried at room temperature, and coverslipped with water-soluble mounting media (Sigma-Aldrich). Digital images were captured from at least three randomly chosen visual fields using the Olympus BX53 microscopy imaging system with the same settings for each section.

### Hepatic glycogen measurement

Hepatic glycogen was measured according to manufacturer’s instructions (SolarBio, Shanghai, China). In brief, 100 mg of liver tissues were homogenized in 0.75 mL of extraction solutions (Reagent I). The homogenates were boiled in a water bath for 20 minutes, gently mixed every 5 minutes. After being cooled down to room temperature, the samples were transferred to a fresh tube containing 5 mL distilled water, mixed thoroughly, and centrifuged at 8000x g for 10 minutes. The resulting supernatants were transferred to a new tube. Subsequently, 60 μL from each sample were transferred into a well of a 96-well plate and mixed with 40 μL of Reagent II. The absorbance at 620 nm was recorded in a microplate reader (Biotech, Synergy LX). Glycogen content was expressed as mg per g live tissues.

### RNA sequencing analysis

ASCs were isolated as previously described (*32*). Briefly, abdominal fat pads were collected aseptically from three WT and three trappc9 KO mice aged 3-4 weeks. Tissues were washed three times in Hank’s balanced salt solutions, minced into small pieces, and then treated with 1 mg mL-1 type-I collagenase at 37 °C with gently mixing at 120 rpm. Enzymatic digestions were terminated by adding 10 mL of α-MEM media containing 10% FBS, 1x L-glutamine, and 1x penicillin/ streptomycin. The digestion mixtures were filtered through a 70-μm cell strainer (ThermoFisher), and the filtrates were centrifuged at 1200 rpm for 5 min. The cell pellets were washed once in phosphate saline and resuspended in 1 mL TRIzol reagents (Invitrogen) for RNA sequencing (RNAseq) analysis.

DRD2-containing neurons were immunomagnetically isolated from both WT and trappc9 KO mice at age 3-4 weeks. In brief, brain tissue was minced into small pieces and treated with 30 U/mL papain at 37 °C for 20 minutes in neurobasal medium supplemented with 1% penicillin-streptomycin, 1% L-glutamine, 2% B27, and 2500 U DNase I. The digestion was terminated by adding 1 mL fetal bovine serum (FBS). The digested tissues were gently dissociated by pipetting through a blue tip and subsequently passed through 70-μm and 40-μm cell strainers (ThermoFisher). Cells in the flowthroughs were collected by a centrifugation at 200 x *g* for 10 minutes and washed once with neurobasal medium with 10% FBS and all supplements as above. Cells were resuspended in PBS (pH 7.2) containing 0.5% bovine serum albumin, 0.5 mM EDTA, 5 μg/mL insulin, and 1 g/L glucose and incubated with DRD2-specific antibodies pre-coupled on magnetic beads at 4 °C for 15 minutes. Cells on beads were washed once in above PBS buffers, separated on a magnetic grate, and resuspended in 1 mL TRIzol reagents for RNAseq analysis.

RNAseq and data processing were performed at Novogene (Tianjin, China). Libraries were prepared using oligo(dT)-attached magnetic beads. Sequencing was performed on the Illumina platform. Raw data underwent through quality control analyses to remove adapters and low-quality reads. The processed reads were aligned to mouse genome (GRCm39 assembly), and gene expression was quantified via Fragments Per Kilobase of transcript per Million mapped reads (FPKM). Differential gene expression was modeled in the DESeq2 R package (1.20.0) or edgeR R package (3.22.5). The resulting p values were adjusted using the Benjamini & Hochberg method. Adjusted p value of 0.05 and two-fold changes were set as the threshold for significantly differential expression.

For enrichment analysis, Gene Ontology (GO) enrichment analysis of differentially expressed genes was implemented by the clusterProfiler R package (3.8.1), in which gene length bias was corrected. GO terms with Benjamini-Hochberg-adjusted p value < 0.05 were considered significantly enriched. The clusterProfiler R package (3.8.1) was applied to test the statistical enrichment of differentially expressed genes in KEGG pathways. Protein-protein interaction networks of neurotensin receptors and dopamine receptors were obtained from the String database (www.string-db.org).

### Synaptosome preparation and proteomic analysis

Synaptosomes were isolated from fresh brain tissues as previously described (*67*). Briefly, WT and trappc9 KO mice aged at 4-5 months were anesthetized. The brain was removed and homogenized with a dounce homogenizer on ice in 7ml freshly prepared homogenization buffer (0.32 M sucrose, 1mM EDTA, 1M DTT, protease inhibitor). Homogenates were centrifuge at 4°C 1000 x g for 10 min. The resulting supernatants were layered onto 1.2M sucrose in 14 x 89 mm Ultra-Clear centrifuge tubes (Beckman, Palo Alto CA). The gradients were centrifuge at 4 °C 160,000 x g for 35 min (SW41 rotor) in a Beckman L8-80 M Ultracentrifuge. On the gradients, synaptosomes were visible as a cloudy band at the interface between 0.32M and 1.2M sucrose and collected. Protein concentrations in synaptosomes were measured.

Label-free quantitative proteomics analysis was conducted as previous described (*68*). In brief, synaptosomes (10μg proteins) from each brain were mixed with an equal volume of acetone pre-cold to -20°C. After incubation for 60 minutes at -20°C, the samples were centrifuged at 4°C 14, 000 x g for 10 minutes. The pellets were dried in air for 30 minutes and subjected to mass spectrometry analysis at the Beth Israel Deaconess Medical Center Mass Spectrometry Core facility (https://www.bidmc.org/research/core-facilities/mass-spectrometry-proteomics-metabolomics-core). The LC-MS/MS spectra of peptides were analyzed using the Mascot Version 2.7.0 (Matrix Science) by searching the reversed and concatenated mouse protein database (Mouse_20221214.fasta) with a parent ion tolerance of 18 ppm and fragment ion tolerance of 0.05 dalton. Raw data of total spectrum count (TSC) were imported into Scaffold (v5.2.2) software (proteome software, Inc) with a peptide threshold of ∼50%, protein threshold of ∼95%, and at least 2 peptides. Further statistical analysis was performed using RStudio (v4.3.1). The raw data were adjusted by adding a value of one to all entries to account for zero counts. Data processing was performed in R studio (v4.3.1) utilizing the limma (v3.58.1) package for linear modeling and differential expression analysis. Differential expression of a protein was assessed using an empirical Bayes method. A p value of 0.05 and a two-fold change were set as the threshold for determining differential expression. GO and KEGG analyses were done as above in RNAseq analysis. Synapse-specific SynGO database was used to retrieve significantly enriched terms describing cellular components.

### SDS-PAGE and Western blot analysis

SDS-PAGE and Western blot analysis were carried out as standard procedures. Primary antibodies used for Western blot analysis included: TH (1:5000, Abcam, ab137869), Unc13-3 (1:1000, Synaptic systems, 126303), Synpo1 (1:2000, Novus biologicals, NBP2- 39100), PSD95 (1:1000, Cell Signaling, 2507S), Dlg2 (1:200, Abcam, ab97290), Nstr2 (1:5000, Novus biologicals, NB-100-56472SS), Gpm6a (1:5000, Synaptic systems, 238003), Insyn1 (1:200, MyBiosource, MBS3218282), GAPDH (1:10000, EMD Millipore, MAB374), and β-actin (1:4000, Sigma-Aldrich, A5441). Peroxidase AffiniPure secondary antibodies (Jackson ImmunoResearch, West Grove, USA) were diluted at 1:10, 000 in PSB containing 2% bovine serum albumin (Sigma-Aldrich). Blots were developed using enhanced Pierce^TM^ enhanced chemiluminescence (ThermoFisher) and imaged with the ChemiDoc^TM^ MP Imaging System (Bio-Rad, USA) or with a CCD camera (Alpha Innotech). Densitometry was conducted using the NIH ImageJ/Fiji software. Background signals were removed. Signal intensities for each protein of interest were normalized with the signal of GAPDH or β-actin in the corresponding samples.

### Immunofluorescent microscopy and image analysis

Mice aged 3 to 5 months were deeply anesthetized and transcardially perfused with 50mL PBS followed by 50mL 4% (w/v) paraformaldehyde (Sigma-Aldrich) in PBS. Fixed brains of WT and trappc9 KO mice were cut into 30 μm thick coronal sections on a cryostat (Leica). Sections were placed in cryoprotectant solutions containing 30% ethylene glycol and 30% glycerol prepared in PBS and stored at -20°C until use.

Immunolabeling was performed with floating brain sections with procedures as previously described (*24, 69*). Sections were removed from cryoprotectant solution and washed three times in PBS at room temperature. Sections were then permeabilized in PBS containing 5% normal donkey serum, 0.2% Triton X-100 and 1% bovine serum albumin for 1 hour and then stained overnight at 4°C with primary antibodies sheep anti-TH (1:1000, Novus biologicals, NB300-109), rabbit anti-Vamp2 (1:1000, Cell signaling technology, #13508), mouse anti-Bassoon (1:400, Enzo life sciences, SAP7F407) and mouse anti-NeuN (1:500, Millipore, MAB377). Afterward, sections were incubated with donkey anti-sheep Alexa 647 (1:500, Jackson immunoresearch, 713-605-147), donkey anti-rabbit Alexa 488 (1:500, Jackson immunoresearch, 711-545-152) and donkey anti-mouse Rhodamine Red-X (1:500, Jackson immunoresearch, 715-295-151) for 2 hours at room temperature. Cells in brain sections were identified by staining with Hoechst34580 (Invitrogen). Brain sections were mounted onto glass slides with ProLong Gold (Invitrogen) for fluorescence microscopy.

Digital images were acquired through a Zeiss LSM 880 confocal microscope (Carl Zeiss, Jena, Germany) with the same settings, including pinhole, laser strength, signal gain and offset. Super-resolution images were captured using 405nm, 488nm, 561nm, and 633nm laser. For quantitative analysis of neurotransmitter release sites, z-stack images across 6 layers with a z-interval of 1.12 μm were captured using a 63x oil immersion objective (Plan-Apochromat, NA1.4) from randomly chosen fields in the NAc. Whole section images were acquired by tile scan using a 10x 0.3 objective and subsequently stitched with ZEN Black software. Composite images were presented as best fit and generated with the Zeiss Zen Blue software. The number of puncta in reconstructed 3-D images was analyzed using the Cell Counter plugin in ImageJ/Fiji software v2.9.0. For analyzing the co-localization of Vamp2, Bassoon and TH, images from 6 randomly chosen visual fields in the NAc were examined per brain section.

## Acknowledgments

We are grateful to John M Asara at the Beth Israel Deaconess Medical Center Mass Spectrometry Core facility for proteomic analysis.

## Author contributions

Y.L., M.U., E.S., Y.K., and A.B. designed, performed, and analyzed experiments. Y.L., M.U., Y.K., and Z.W. provided histological evaluations and data analysis. Y.L., M.D., and X.L. wrote the manuscript. X.L. conceived, designed, analyzed, and supervised the overall direction of the study.

## Data and materials availability

All data needed to evaluate the conclusions in the paper are present in the paper and/or the Supplementary Materials. Additional data related to this paper may be requested from the authors.

## Supplementary figures and figure legends

**Figure S1.**
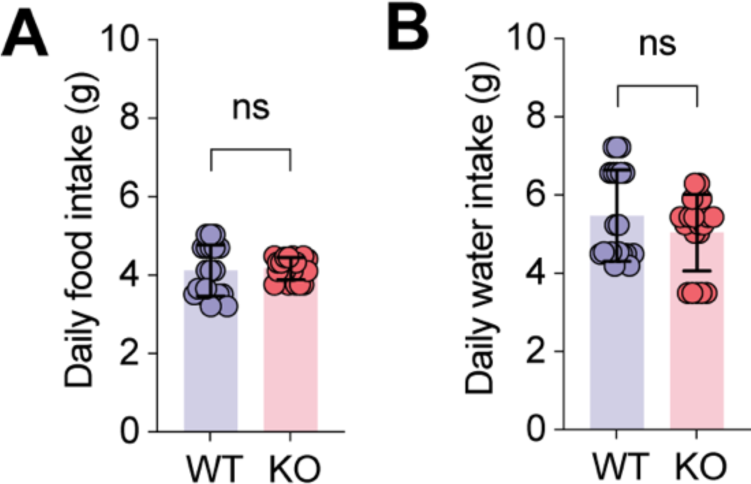
The abrogation of trappc9 in mice does not alter water and food intake. Daily food (**A**) and water (**B**) intake of WT and trappc9 mice were measured for one week at age 5 months (N=12 mice per genotype). Each symbol in bar graphs represents one mouse. Data are Mean±SD. Two-tailed Student’s t-test: ns, no significance.

**Figure S2.**
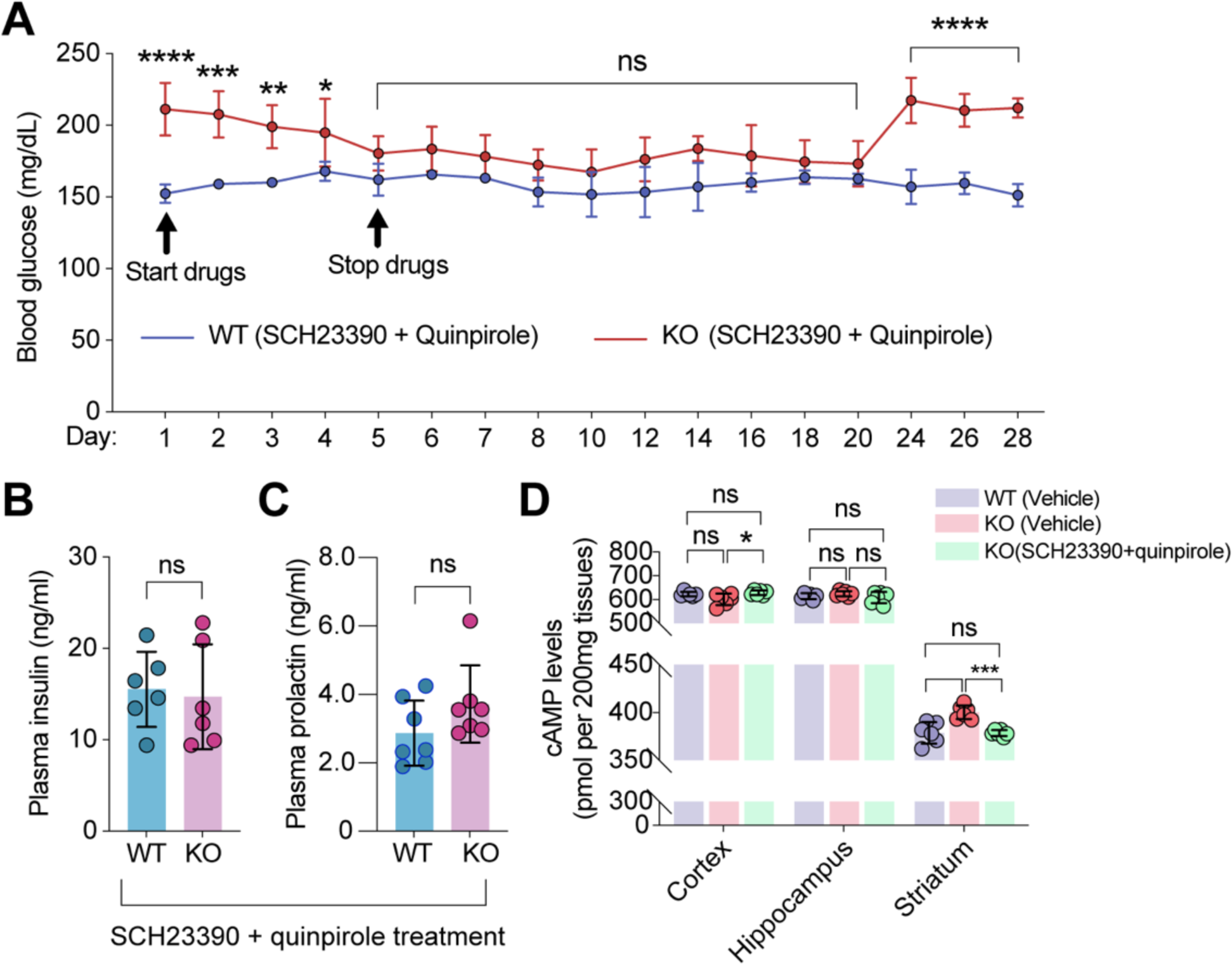
The combined treatment with SCH23390 and quinpirole restores the homeostasis of systemic glucose, insulin and prolactin, and signs of their actions in the brain. A) Efficacy of the SCH23390 and quinpirole combination in lowering blood glucose in trappc9 KO mice. The plot shows that daily administration of both drugs had a trivial effect on blood glucose levels in WT mice, but effectively decreased blood glucose levels in trappc9 KO mice 5 days after treatment. Drug effects lasted for two weeks after the treatment was stopped. Efficacy of the combined treatment in correcting hyperinsulinemia (**B**) and hyperprolactinemia (**C**) in trappc9 KO mice. **D**) Effects of the treatment combing SCH23390 and quinpirole on cAMP levels in the indicated brain areas. Efficacy studies in (**A**) were done with a cohort of 7 mice for each genotype. Each symbol in bar graphs represents one mouse. Data are Mean±SD. Statistical significance was determined by two-tailed Student’s t-test (**A**, **B**, **C**) or one-way ANOVA and post hoc Tukey’s test (**D**): * P<0.05; ** P<0.01; *** P<0.005; **** P<0.001; ns, no significance.

